# A Spatiotemporal Map of Reading Aloud

**DOI:** 10.1101/2021.05.23.445307

**Authors:** Oscar Woolnough, Cristian Donos, Aidan Curtis, Patrick S. Rollo, Zachary J. Roccaforte, Stanislas Dehaene, Simon Fischer-Baum, Nitin Tandon

## Abstract

Reading words aloud is a fundamental aspect of literacy. The rapid rate at which multiple distributed neural substrates are engaged in this process can only be probed via techniques with high spatiotemporal resolution. We probed this with direct intracranial recordings covering most of the left hemisphere in 46 humans as they read aloud regular, exception and pseudo-words. We used this to create a spatiotemporal map of word processing and to derive how broadband gamma activity varies with multiple word attributes critical to reading speed: lexicality, word frequency and orthographic neighborhood. We found that lexicality is encoded earliest in mid-fusiform (mFus) cortex and precentral sulcus, and is represented reliably enough to allow single-trial lexicality decoding. Word frequency is first represented in mFus and later in the inferior frontal gyrus (IFG) and inferior parietal sulcus (IPS), while orthographic neighborhood sensitivity resides solely in IPS. We thus isolate the neural correlates of the distributed reading network involving mFus, IFG, IPS, precentral sulcus and motor cortex and provide direct evidence for dual-route models of reading, with parallel processes via the lexical route from mFus to IFG, and the sub-lexical route from IPS and precentral sulcus to anterior IFG.

## Introduction

Reading a word aloud depends on multiple complex cerebral computations: mapping the visual input of a letter string to an internal sequence of sound representations, and then their expression through orofacial motor articulations. Models of how this mapping occurs have postulated a dual-route architecture (Coltheart et al., 2001; Perry et al., 2007, 2010, 2019; Taylor et al., 2013), with a lexico-semantic route for rapidly reading known words and a sub-lexical route for constructing phonology of novel words. Contrasts between phonological exception words and pseudowords (Fiebach et al., 2002; Sebastian et al., 2014; Shim et al., 2012; Taylor et al., 2013) are often used to maximally separate these two routes. Exception words contain irregular grapheme-phoneme associations (e.g. yacht, sew) and their pronunciations must therefore be retrieved from internal lexical representations, as they cannot be accurately constructed *de novo*. In contrast, pseudowords have no stored representations and their phonology must be constructed rather than retrieved. Finally, regular words may be read correctly by either route, with the relative speeds of the routes depending on the level of automatization of reading as well as the size of the reader’s vocabulary.

At the brain level, the two routes are thought to share an initial stage of visual word form processing in left occipitotemporal cortex (Dehaene and Cohen, 2011), with only small topological changes depending on whether the task favors lexical or phonological processing (Bouhali et al., 2019). More anteriorly, ventral temporal cortex is strongly implicated as mediating the lexical route, with mid-fusiform cortex (mFus) functioning as the orthographic lexicon, a region where familiar letter strings are mapped onto known words (Glezer et al., 2015; Hirshorn et al., 2016; Kronbichler et al., 2004; Liu et al., 2021; Lochy et al., 2018; Nobre et al., 1994; White et al., 2019; Woolnough et al., 2021). mFus is sensitive to lexicality and word frequency (Kronbichler et al., 2004; White et al., 2019; Woolnough et al., 2021), and its activity is modulated by visual word learning (Glezer et al., 2015; Taylor et al., 2019). Conversely, the sub-lexical route, essential for articulating novel words, is thought to engage the inferior parietal lobe (IPL). The dysfunction of IPL is associated with specific forms of reading deficits, most prominently impairing pseudoword reading (Dickens et al., 2019; Rapp et al., 2016; Raschle et al., 2011; Temple et al., 2003; Tomasino et al., 2020), in addition to broader phonological and semantic deficits (Binder et al., 2009; Hula et al., 2020; Numssen et al., 2021). The two routes of reading are presumed to be active in parallel (Simos et al., 2002) and converge in the inferior frontal gyrus (IFG) (Taylor et al., 2013).

The theoretical embedding of the routes of reading in neural architecture has primarily been derived from lesion data and functional MRI (Bouhali et al., 2019; Jobard et al., 2003; Ripamonti et al., 2014; Tomasino et al., 2020), but these modalities lack the spatiotemporal resolution to delineate the dynamics of these networks and their real-time interactions that allow us to rapidly read. To overcome this, we utilized intracranial recordings in a large cohort of patients (46 patients, 3,846 electrodes), with medically intractable epilepsy, while they read aloud known and novel words, creating a comprehensive 4D map of the spatiotemporal dynamics of behaviorally important lexical and sub-lexical processes throughout the reading system.

## Results

Participants read aloud phonologically regular words, exception words and novel pseudowords (Figure 1A) while concurrent recordings were performed using 3,846 separate intracranial electrodes placed for the localization of intractable epilepsy (Figure 1B,C). 42 participants had depth recordings using stereotactic EEG electrodes (sEEGs) and 4 had subdural grid electrodes (SDEs).

**Figure 1:**
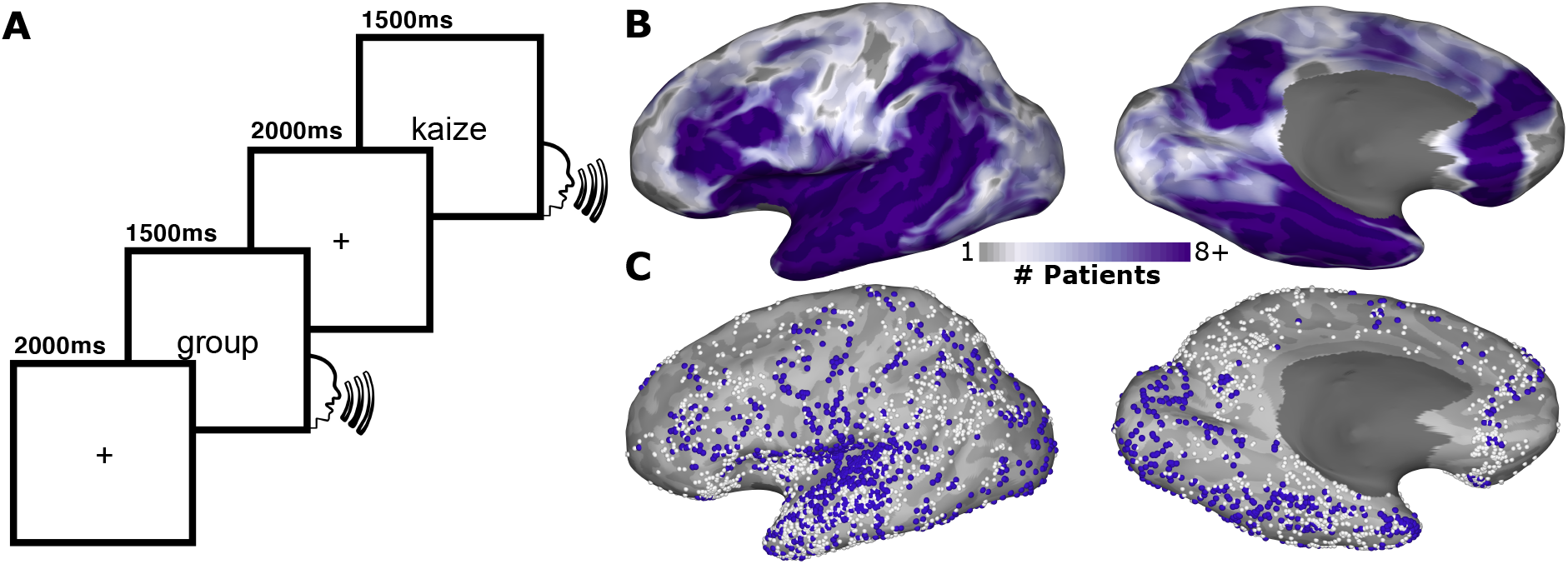
Experimental Design and Electrode Coverage. (A) Schematic representation of the reading task. (B) Representative coverage map (46 patients) and (C) individual electrode locations (3,846 electrodes) for the left hemisphere, highlighting responsive electrodes (1,248 electrodes; >20% activation above baseline).

### Behavioral Analysis

Mean (± SD) response times (RTs) were: regular words (734 ± 113 ms), exception words (738 ± 113 ms) and pseudowords (911 ± 162 ms) (Figure 2A). Regular and exception words showed no difference in RT (Wilcoxon sign rank, p = 0.90; ln(Bayes Factor (BF_10_)) = −1.6) though pseudoword RT was slower than for exception words (p < 10^−8^, ln(BF_10_) = 32). 95 ± 4% of trials were correctly articulated. The most common errors were regularization of exception words (e.g. sew as *sue*, soot as *sute*) or lexicalization of pseudowords (e.g. shret as *shirt*, jinje as *jingle*). Error trials were excluded from analysis.

**Figure 2:**
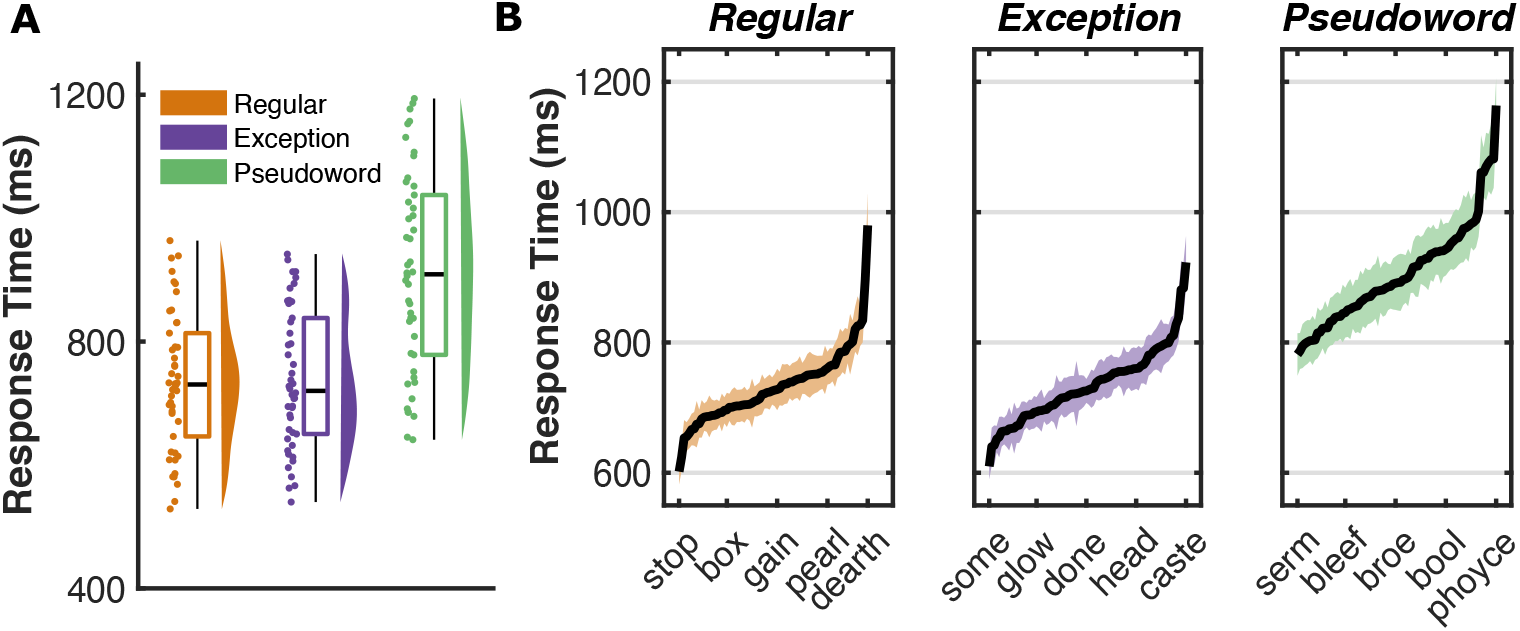
Population Word Response Times. (A) Response time distribution for each of the three word classes, averaged within participant, (B) Mean (± SE) response times for each item within the three word classes, averaged across participants.

To determine the word features that modulate RT, we performed linear mixed effects (LME) and Bayes factor (BF) analyses on each word class with fixed effects modelling linguistic factors commonly linked to word identification and articulation (Table 1). Regular and exception word RTs showed greatest modulation by word frequency. Exception words also displayed modulation by orthographic neighborhood and we observed a significant interaction between regularity and orthographic neighborhood (LME: t(6,589) = −4.2, β = −62.5, p < 10^−4^, 95% CI −91.7 to −33.3). Pseudoword RT was most strongly associated with orthographic neighborhood, meaning pseudowords with many known word neighbors were articulated faster.

**Table 1:**
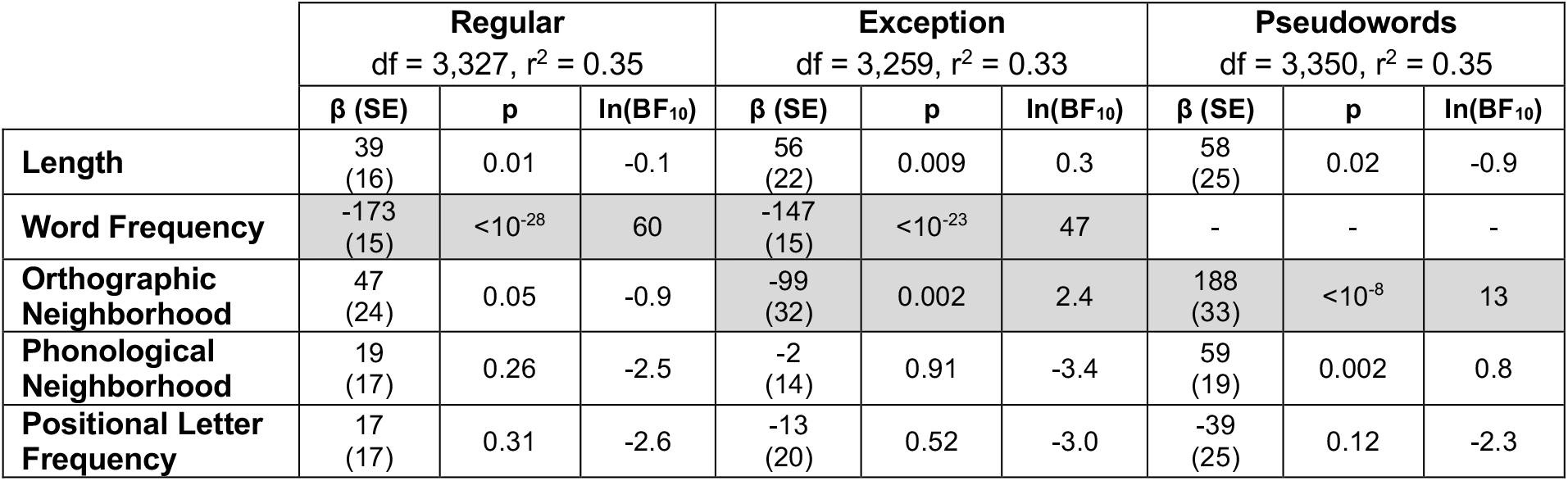
Statistical Modelling of Response Time. As predictors were normalized, β values approximate change in RT between extreme values within the entire stimulus set (Supplementary Table 1). Factors with strong evidence of an effect (ln(BF_10_) > 2.3) are highlighted.

| | Regular<br>df = 3,327, $r^2 = 0.35$ | | | Exception<br>df = 3,259, $r^2 = 0.33$ | | | Pseudowords<br>df = 3,350, $r^2 = 0.35$ | | |
| --- | --- | --- | --- | --- | --- | --- | --- | --- | --- |
| | $\beta$ (SE) | p | $\ln(\text{BF}_{10})$ | $\beta$ (SE) | p | $\ln(\text{BF}_{10})$ | $\beta$ (SE) | p | $\ln(\text{BF}_{10})$ |
| **Length** | 39<br>(16) | 0.01 | -0.1 | 56<br>(22) | 0.009 | 0.3 | 58<br>(25) | 0.02 | -0.9 |
| **Word Frequency** | -173<br>(15) | $<10^{-28}$ | 60 | -147<br>(15) | $<10^{-23}$ | 47 | - | - | - |
| **Orthographic Neighborhood** | 47<br>(24) | 0.05 | -0.9 | -99<br>(32) | 0.002 | 2.4 | 188<br>(33) | $<10^{-8}$ | 13 |
| **Phonological Neighborhood** | 19<br>(17) | 0.26 | -2.5 | -2<br>(14) | 0.91 | -3.4 | 59<br>(19) | 0.002 | 0.8 |
| **Positional Letter Frequency** | 17<br>(17) | 0.31 | -2.6 | -13<br>(20) | 0.52 | -3.0 | -39<br>(25) | 0.12 | -2.3 |

### Spatiotemporal Mapping of Single Word Reading

We used a mixed-effects, multilevel analysis (MEMA) of broadband gamma activity (BGA; 70-150 Hz) in group surface normalized space to create a population level map of cortical activation across the population. This analysis is specifically designed to account for sampling variations and to minimize effects of outliers (Argall et al., 2006; Conner et al., 2014; Esposito et al., 2013; Fischl et al., 1999; Kadipasaoglu et al., 2014; Saad and Reynolds, 2012). All correctly articulated trials were used. A 4D representation of activation on the cortical surface was generated by collating MEMA on short, overlapping time windows (150 ms width, 10 ms spacing) to generate successive images of cortical activity, time locked to stimulus onset (Video 1) or the onset of articulation (Video 2). The spatial distribution of activations was highly consistent across word classes (Supplementary Figure 1).

By collapsing across these frames, we visualized peak activations at each point on the cortical surface (Figure 3A). Further, to create a more focused visualization of the spatiotemporal progression across reading-sensitive cortex, we selected regions of interest (ROIs) at sites with maximal activation, that also corresponded to sites believed to be important for written word processing, speech production and speech monitoring (Figure 3B,C, Supplementary Figure 2).

**Figure 3:**
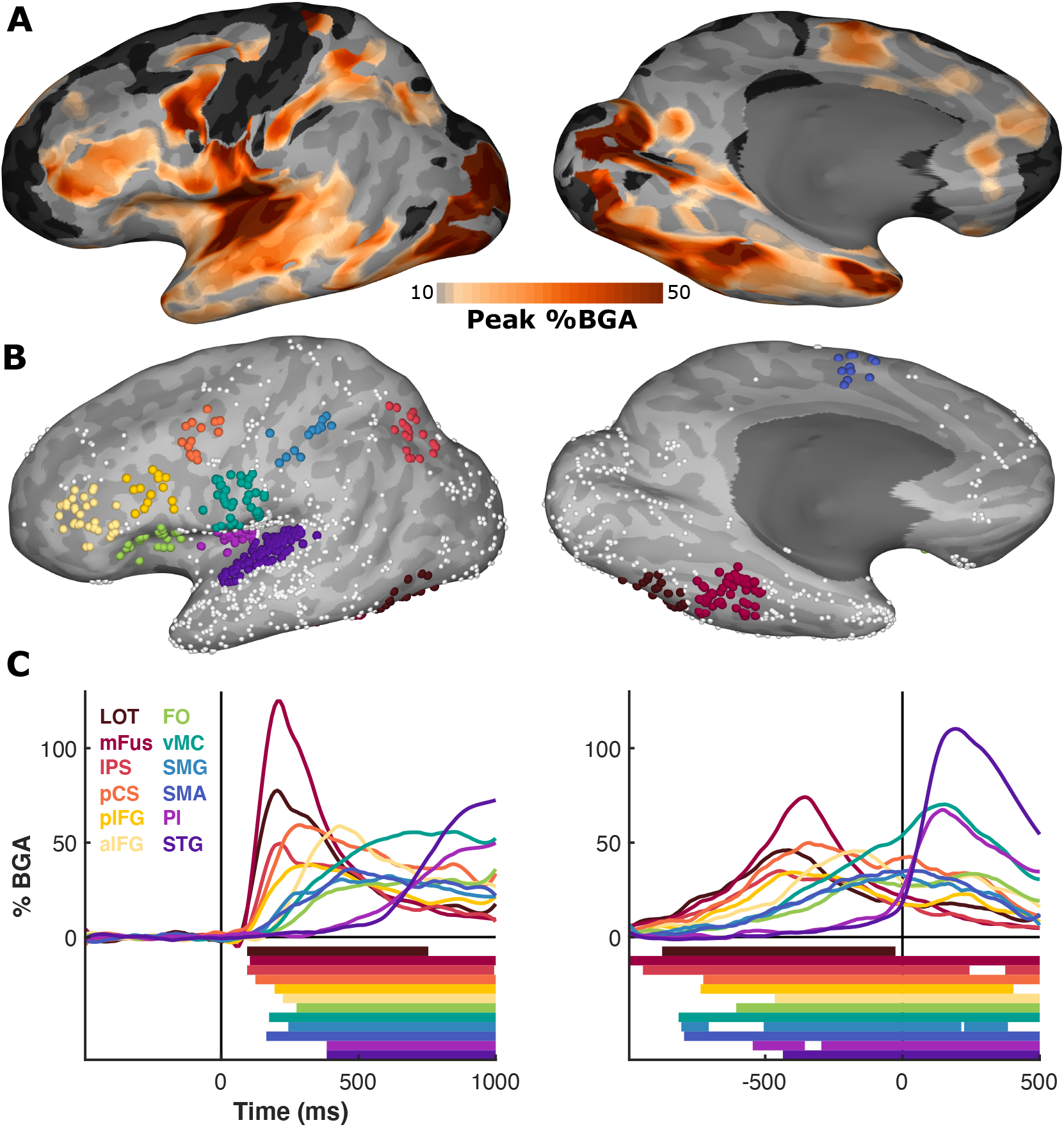
Spatiotemporal Profile of Cortical Activations. (A) Collapsed articulation-locked activation movie (Video 2) highlighting the amplitude of peak activation. (B) Representative ROIs in 12 anatomically and functionally distinct regions, showing all responsive electrodes. (C) Mean activation during word reading of each ROI, averaged within patient, time locked to stimulus onset (left) and articulation onset (right). Standard errors omitted for visual clarity. Colored bars represent regions of activation greater than baseline (Wilcoxon signed-rank, q < 0.05). LOT, Lateral OccipitoTemporal cortex; mFus, mid-Fusiform Cortex; IPS, Inferior Parietal Sulcus; pCS, pre-Central Sulcus; pIFG, posterior Inferior Frontal Gyrus; aIFG, anterior Inferior Frontal Gyrus; FO, Frontal Operculum; vMC, ventral Motor Cortex; SMG, Supra Marginal Gyrus; SMA, Supplementary Motor Area; PI, Posterior Insula; STG, Superior Temporal Gyrus.

### Spatiotemporal Representation of Lexical Factors

To distinguish activity patterns across word classes we contrasted grouped gamma power activations between exception vs. pseudowords (lexicality) and exception vs. regular words (regularity). The lexicality contrasts demonstrated clusters in early visual cortex, mFus, precentral sulcus (pCS), inferior parietal sulcus (IPS) and anterior inferior frontal gyrus (aIFG). The regularity contrast only demonstrated a small cluster in pCS.

To distinguish whole-brain activity patterns for each of these factors, within-individuals at a single trial level, we used a logistic regression decoder. Decoders trained to distinguish between exception word and pseudoword trials demonstrated high decoding accuracy, with some patients showing >80% decoding accuracy (Figure 4C). These lexicality decoders displayed high electrode weightings in the same regions as the lexicality contrasts listed above with the addition of the superior frontal sulcus (SFS) and the ventral visual stream – thus an independent analysis corroborates the critical role of the LOT, mFus, pCS, IPS and aIFG in lexicality processing (Figure 4D). Decoders trained to distinguish exception and regular words did not show significantly greater decoding accuracy than in the baseline period.

**Figure 4:**
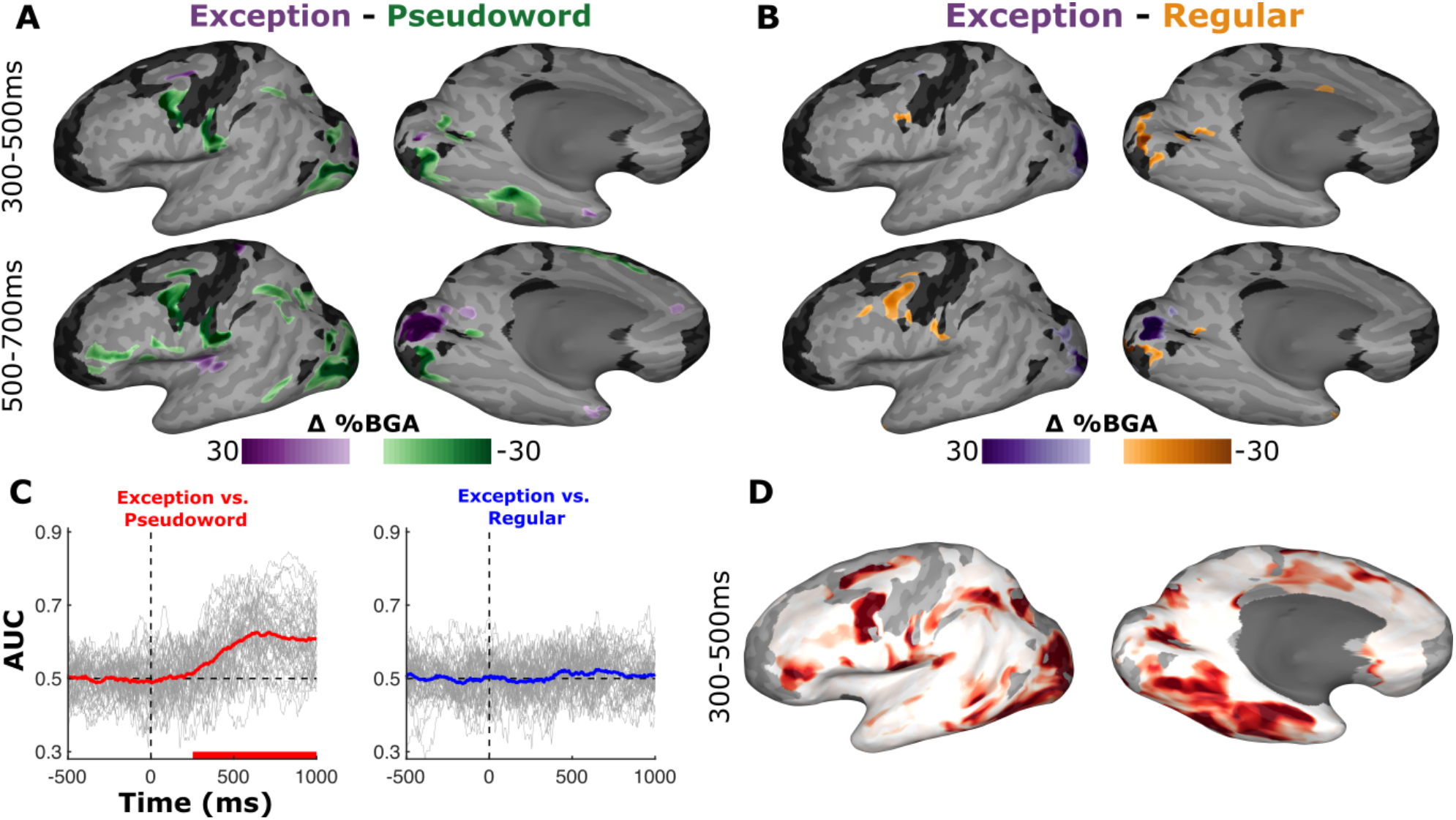
Contrasting Word Classes. (A,B) MEMA contrasts of (A) exception – pseudoword and (B) exception – regular, revealing regions of significantly different BGA between conditions (p < 0.01 corrected). Regions in black did not have consistent coverage for reliable MEMA results. (C) Decoding accuracies of the logistic regression decoders trained to distinguish exception word vs. pseudoword trials (left) and exception word vs regular word trials (right). Grey lines represent individual patient decoding accuracies. Colored line represents median accuracy. Colored bars represent time periods significantly greater than chance (Wilcoxon signed-rank, q < 0.05). (D) Cortical surface representation of population average electrode weightings of the exception vs pseudoword decoder between 300 – 500 ms.

To characterize the timing of lexicality distinctions between known words (regular and exception) and novel pseudowords broadly across ROIs (Figure 5), we compared their activity by word class. Lexicality distinctions, with stronger activity for pseudowords, were observed earliest in mFus and subsequently in IFG and IPS. Distinctions were also observed in the opposite direction in post-articulatory auditory regions (posterior insula and superior temporal gyrus) relating to differences in RT between known and novel words.

**Figure 5:**
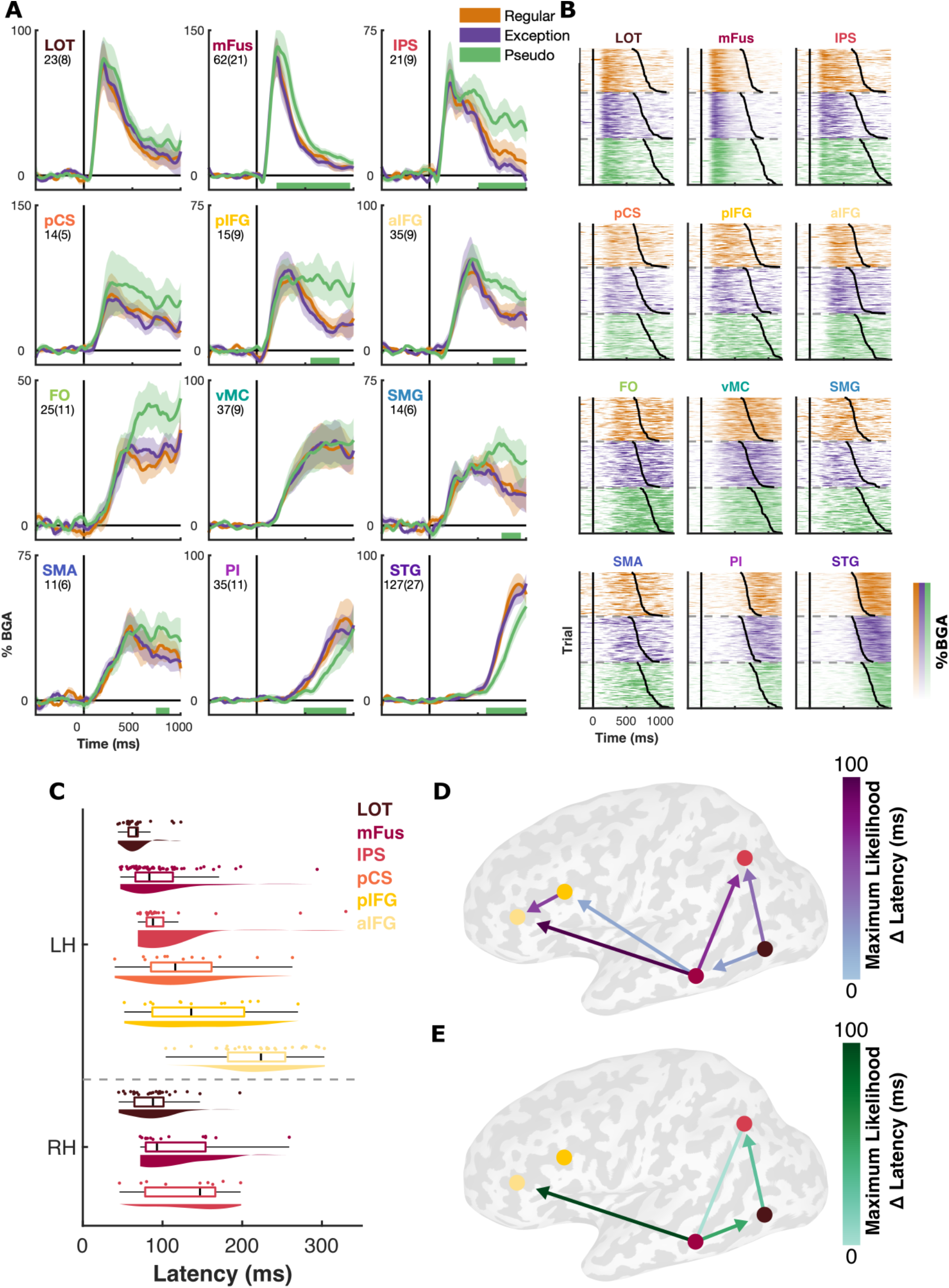
Spatiotemporal Activation Profiles of Known and Novel Words. (A) Mean activation (± SE) for each word class, within each ROI, during word reading, averaged within patient, time locked to stimulus onset (See also Supplementary Figure 2 for right hemisphere ROIs). Number of electrodes and patients, per ROI, is indicated. Colored bars represent regions of significant difference from exception words (Wilcoxon signed-rank, p < 0.05 for >100ms). (B) Mean activation per unique word, within each ROI. Trials separated by word class and sorted by the word’s mean RT for the patients included within the given ROI. Mean RT for each word is highlighted. (C) Latency of first onset of activation (first derivative of BGA >3.5SD above baseline) for each electrode within each ROI. (D,E) Network representations demonstrating maximum likelihood latency differences between ROIs, within patients with simultaneous coverage, for (D) initial activation latency and (E) initial lexicality distinction latency. Lines excluded for ROI pairs where there was no simultaneous coverage of significant electrodes. Arrowheads not shown for differences <10ms. LOT, Lateral OccipitoTemporal cortex; mFus, mid-Fusiform Cortex; IPS, Inferior Parietal Sulcus; pCS, pre-Central Sulcus; pIFG, posterior Inferior Frontal Gyrus; aIFG, anterior Inferior Frontal Gyrus; FO, Frontal Operculum; vMC, ventral Motor Cortex; SMG, Supra Marginal Gyrus; SMA, Supplementary Motor Area; PI, Posterior Insula; STG, Superior Temporal Gyrus.

Within patients that had simultaneous electrode coverage within multiple ROIs, we compared onset latency of activation, across all word classes (Figure 5D), and the initial latency of lexicality distinction (Figure 5E). This demonstrated the spread of initial activation from LOT to mFus and IPS before spreading to pIFG then aIFG. In contrast the earliest lexicality distinctions were observed in mFus which subsequently spread to LOT and aIFG. None of the individual tested pIFG electrodes showed a significant lexicality distinction.

For the six ROIs that showed a clear pre-articulatory peak in activation, we analyzed their activity for sensitivity to the main drivers of RT seen in the behavioral analysis; lexicality, word frequency of known words and orthographic neighborhood for pseudowords. mFus showed the earliest sensitivity to lexicality, followed by LOT and pCS, and then broad sensitivity across multiple regions (Figure 6A). mFus showed an early and long-lasting word frequency sensitivity, with IPS and aIFG becoming sensitive later (500-700 ms). Sensitivity to orthographic neighborhood of pseudowords was only seen in IPS (500-700 ms). In the right hemisphere we observed effects of lexicality only in LOT and IPS (with no involvement of right mFus) and no effect of word frequency or orthographic neighborhood. (Supplementary Figure 2).

**Figure 6:**
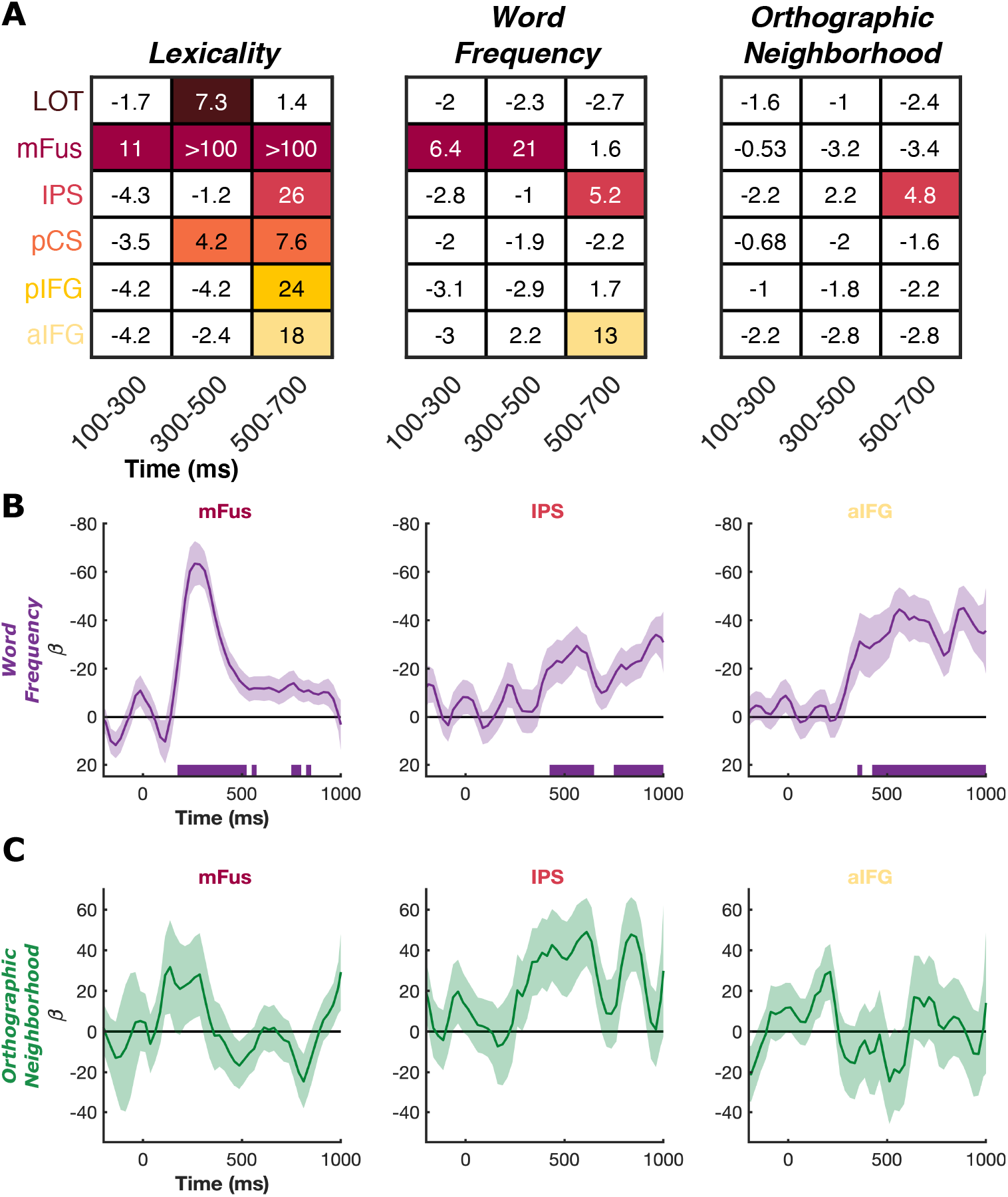
Regression of Lexical Factors. (A) Bayes factor analysis of lexicality, word frequency and orthographic neighborhood effects in the six pre-articulatory ROIs, for three time windows. Lexicality tested all known words against pseudowords. Word frequency was regressed across all known words. Orthographic neighborhood was regressed across all pseudowords. Bayes factor (ln(BF_10_)) shown for each contrast and values >2.3 are highlighted. (B,C) Linear mixed effects (LME) model regression of (B) word frequency in known words and (C) orthographic neighborhood in pseudowords, in three ROIs (β ± SE; mFus, 62 electrodes, 21 patients; IPS, 21 electrodes, 9 patients; aIFG, 35 electrodes, 9 patients). Colored bars represent regions of significance (q < 0.05).

For the three regions with evidence of word frequency or orthographic neighborhood effects, mFus, IPS and aIFG, we generated LME models at a higher time resolution. Sensitivity to word frequency was observed earliest in mFus (200 ms) followed by IPS and aIFG (425 ms) (Figure 6B). In IPS we observed a period of elevated orthographic neighborhood sensitivity, but this did not show significance at this time resolution (Figure 6C).

## Discussion

This work comprehensively maps the spatiotemporal spread of cortical activation across the left hemisphere during word reading to derive the dynamics of cortical networks underlying literacy, using intracranial recordings in a large population. We find two regions specifically selective for lexicality - the mFus and pCS, with subsequent engagement of the IPS and IFG. This lexicality network is broadly comparable with the spatial substrates derived from fMRI (Heim et al., 2013; Taylor et al., 2013, 2014), with an added benefit of millisecond temporal resolution. Responses in lexicality sensitive regions maximally separate for known words vs pseudowords between 300-500ms after stimulus onset, in a manner reliable enough to enable above-chance single-trial decoding of lexicality. These data minimize the impact of response time variation, which confounds modalities with lower temporal resolution (e.g. fMRI) and may artificially inflate lexicality effects in regions such as IFG (Taylor et al., 2014). Our findings corroborate the theoretical dual-route architecture of reading (Coltheart et al., 2001; Perry et al., 2007, 2010, 2019), primarily derived from behavioral and neuropsychological data.

We have previously demonstrated that mFus is the earliest region in ventral temporal cortex to show sensitivity to word frequency while reading (Woolnough et al., 2021). It is often assumed that sensitivities to statistical properties of language, such as word frequency, seen in ventral temporal cortex arise solely from a top-down modulation from IFG (Heim et al., 2013; Liu et al., 2021; Price and Devlin, 2011; Whaley et al., 2016; Woodhead et al., 2014). Here, we demonstrate again the primacy of mFus in coding both word frequency and lexicality, preceding the engagement of aIFG and IPS in these processes by over 200 ms. This suggests that the directional connectivity metrics derived in these studies may not necessarily detect the transmission of lexical information, but perhaps higher-order semantic or phonological information. Additionally, we observe that right LOT demonstrates lexical sensitivity at a comparable time to left LOT, subsequent to the lexicality distinction in left mFus. This likely indicates left mFus provides top-down influences to both ipsilateral and contralateral visual areas. These results consolidate the role of mFus as a specialized orthographic lexicon, organized based on statistical regularities of individual words in natural language.

The IPS was the only region with sensitivity to the orthographic neighborhood of novel pseudowords. IPS has previously been implicated in the grapheme-phoneme conversion process (Dehaene-Lambertz et al., 2018; Xu et al., 2020). Given that IPS shows both word frequency and lexicality sensitivity, its role in sub-lexical processing might appear to be questionable. However, it should be remembered that both routes are thought to activate in parallel and compete for speed (Simos et al., 2002). For known words of high enough frequency, the lexical route is faster and more accurate than the sub-lexical route – thus, once a letter string is identified as a known lexical object, sub-lexical processes are no longer required (unless the word is visually degraded, thus promoting a return to slow left-to-right analysis (Cohen et al., 2008; Vinckier et al., 2006)). Given that the speed of lexical identification varies with word frequency, the timing of the cessation of sub-lexical processes should also be frequency dependent. This interpretation is entirely consistent with our data as IPS shows more sustained activity, but not higher peak activity for pseudowords compared to words. We also observed lexicality effects in right IPS, consistent with TMS studies demonstrating the importance of right, non-dominant parietal cortex in phonological (Hartwigsen et al., 2010) and visuospatial processing (Cazzoli et al., 2015) during reading.

It is theorized that pCS is involved in articulatory phonological processing, specifically feedforward control of articulator velocity (Matchin and Hickok, 2020; Tourville and Guenther, 2011). Through lesion studies, pCS has also been linked to phonological dyslexia (Rapcsak et al., 2009; Tomasino et al., 2020). Our data demonstrate that pCS activation begins early, preceding the IFG, suggesting a role in early linguistic or phonological processing, potentially as part of the sub-lexical route. This is concordant with recent findings suggesting pCS is involved in the grapheme-to-phoneme conversion process (Kaestner et al., 2021). pCS demonstrates lexical sensitivity but no effect of word frequency or orthographic neighborhood. Given the association of pCS with articulation phonology and phonological dyslexia, pCS may contribute to the process of constructing articulatory representations for both words and pseudowords, likely in a manner distinct from the processing in IPS.

This study provides further evidence that medial frontal operculum is involved in pre-articulatory, preparatory processes, distinct from those of the lateral IFG (Mălîia et al., 2018; Woolnough et al., 2019). Lesions involving this region have been linked to impairment of complex articulation (Baldo et al., 2011) which may explain the greater engagement during pseudoword articulation.

We observed no substantial pre-articulatory activity in angular gyrus, instead peaking prominently after the initiation of articulation. This region has previously been linked to semantic and phonological processes during word processing (Graves et al., 2010; Hartwigsen et al., 2010; Sliwinska et al., 2015; Stoeckel et al., 2009). This region appears to be engaged during reading in children but may not be recruited in adults for simple reading tasks (Martin et al., 2015), instead being used primarily for comprehending multi-word phrases (Dronkers et al., 2004; Fridriksson et al., 2018; Matchin et al., 2017; Wilson et al., 2014).

## Conclusions

This dataset comprehensively maps the dynamic functional characteristics of the two reading routes. Extant models suggest that all words should be initially processed through both reading pathways, with areas responsible for lexical access being activated inversely proportionally to lexical frequency, as demonstrated here in mFus and aIFG. Once a known word is detected, the sub-lexical route is interrupted (with activity returning to baseline level), and the word is named through the lexical route. However, if the word is novel, there are two possibilities: (1) that the lexical route is interrupted, with activity returning to baseline or (2) that the lexical route continues to attempt to identify the pseudoword while the sub-lexical route constructs the phonology of these words. Our data clearly implicate the second scenario and provide evidence that the activity in the sub-lexical pathway, through the IPS is driven by pseudoword orthographic complexity. Lastly, the early, lexically sensitive activation of ventral premotor cortex (pCS) implies a role in early grapheme-to-phoneme conversion, potentially as part of the sub-lexical route, distinct from the IPS.

## Materials and Methods

### Participants

46 patients (26 male, 19-60 years, 5 left-handed, IQ 94 ± 14, Age of Epilepsy Onset 18 ± 10 years) participated in the experiments after giving written informed consent. All participants were semi-chronically implanted with intracranial electrodes for seizure localization of pharmaco-resistant epilepsy. Participants were excluded if they had confirmed right-hemisphere language dominance or a significant additional neurological history (e.g. previous resections, MR imaging abnormalities such as malformations or hypoplasia). One additional participant was tested but excluded from analysis as their response times were considered outliers for all tested word classes. All experimental procedures were reviewed and approved by the Committee for the Protection of Human Subjects (CPHS) of the University of Texas Health Science Center at Houston as Protocol Number HSC-MS-06-0385.

### Electrode Implantation and Data Recording

Data were acquired from either subdural grid electrodes (SDEs; 4 patients) or stereotactically placed depth electrodes (sEEGs; 42 patients). SDEs were subdural platinum-iridium electrodes embedded in a silicone elastomer sheet (PMT Corporation; top-hat design; 3mm diameter cortical contact), and were surgically implanted via a craniotomy (Pieters et al., 2013; Tandon, 2012; Tong et al., 2020). sEEGs were implanted using a Robotic Surgical Assistant (ROSA; Medtech, Montpellier, France) (Rollo et al., 2020; Tandon et al., 2019). Each sEEG probe (PMT corporation, Chanhassen, Minnesota) was 0.8 mm in diameter and had 8-16 electrode contacts. Each contact was a platinum-iridium cylinder, 2.0 mm in length and separated from the adjacent contact by 1.5 - 2.43 mm. Each patient had 12-20 such probes implanted. Following surgical implantation, electrodes were localized by co-registration of pre-operative anatomical 3T MRI and post-operative CT scans in AFNI (Cox, 1996). Electrode positions were projected onto a cortical surface model generated in FreeSurfer (Dale et al., 1999), and displayed on the cortical surface model for visualization (Pieters et al., 2013). Intracranial data were collected during research experiments starting on the first day after electrode implantation for sEEGs and two days after implantation for SDEs. Data were digitized at 2 kHz using the NeuroPort recording system (Blackrock Microsystems, Salt Lake City, Utah), imported into Matlab, initially referenced to the white matter channel used as a reference for the clinical acquisition system and visually inspected for line noise, artifacts and epileptic activity. Electrodes with excessive line noise or localized to sites of seizure onset were excluded. Each electrode was re-referenced to the common average of the remaining channels. Trials contaminated by inter-ictal epileptic spikes were discarded.

### Stimuli and Experimental Design

All patients undertook task reading aloud single words and pseudowords. Stimuli were presented on a 2,880 × 1,800 pixel, 15.4” LCD screen positioned at eye-level, 2-3’ from the patient. Participants were presented with 80 each of monosyllabic (i) phonologically regular words, (ii) phonologically irregular exception words and (iii) novel pseudowords and asked to read them aloud. Stimuli were presented using Psychophysics Toolbox (Kleiner et al., 2007) in Matlab, in all lower-case letters, in Arial font with a height of 150 pixels (∼2.2° visual angle). Each stimulus was displayed for 1,500 ms with an inter-stimulus interval of 2,000 ms. Stimuli were presented in two recording sessions, each containing presentation of 120 stimuli in a pseudorandom order with no repeats.

### Signal Analysis

Analyses were performed by first bandpass filtering raw data of each electrode into broadband gamma activity (BGA; 70-150Hz) following removal of line noise (zero-phase 2nd order Butterworth bandstop filters). A frequency domain bandpass Hilbert transform (paired sigmoid flanks with half-width 1.5 Hz) was applied and the analytic amplitude was smoothed (Savitzky - Golay finite impulse response, 3rd order, frame length of 201 ms). BGA is presented here as percentage change from baseline level, defined as the period −500 to −100 ms before each word presentation.

Electrodes were tested to see if they were responsive during the task. Responsiveness was defined as displaying >20% average BGA over baseline for at least one of the three following windows: 100 to 500 ms following stimulus onset, −500 to −100 ms before articulation onset or 100 to 500 ms following articulation onset. Of the 3,846 useable electrodes, 1,248 electrodes were designated responsive based on these criteria.

ROIs were selected in canonical language areas, areas identified in previous intracranial studies of reading (Hirshorn et al., 2016; Lochy et al., 2018; Woolnough et al., 2019, 2021) and areas of high peak activation in the word-type independent MEMA movie. ROI centers were defined on the cortical surface and all responsive electrodes within a set geodesic radius of this point were included. This method was selected as currently available cortical surface parcellations have been shown to be inadequate at predicting functional boundaries of task-evoked activity (Zhi et al., 2021). An exception was made for the STG ROI which was defined using a Human Connectome Project derived parcellation (A1, A4, PBelt, MBelt, LBelt) (Glasser et al., 2016).

Frequentist statistical methods were corrected for multiple comparisons using a Benjamini-Hochberg false detection rate (FDR) threshold of q < 0.05.

### Neural Decoding

Decoding analyses were performed using logistic regression classifiers, using 5-fold cross validation, implemented within MNE-Python (Gramfort, 2013; Gramfort et al., 2014). For each patient, decoding performance was summarized with an area under the curve (AUC) and a set of classifier weights for each electrode. Temporal decoding was performed on BGA using a sliding estimator at each time point, using all available electrodes. Spatial distribution of classifier weights was reconstructed by a cortical surface transform onto a standardized brain surface using each electrode’s presumed “recording zone”, an exponentially decaying geodesic radius (Kadipasaoglu et al., 2014). Cortical surface maps were amplitude normalized within patient then averaged across patient to create a population weighting map.

### Linguistic Analysis

We quantified word frequency as the base-10 log of the SUBTLEXus frequency (Brysbaert and New, 2009). This resulted in a frequency of 1 meaning 10 instances per million words and 4 meaning 10,000 instances per million words. There was no significant difference between word frequency of regular (1.5 ± 0.35; Mean ± SD) and exception (1.7 ± 1.0) words (Wilcoxon rank sum, p = 0.36) (Supplementary Table 1). Positional letter frequency was calculated as the base-10 log of the sum of the SUBTLEXus frequencies of all words with a given letter in a specific ordinal position. Orthographic neighborhood was quantified as the orthographic Levenshtein distance (OLD20); the mean number of single character edits required to convert the word into its 20 nearest neighbors with a log frequency greater than 0 (Yarkoni et al., 2008). Phonological neighborhood densities were obtained from the Irvine Phonotactic Online Dictionary (IPhOD) (Vaden et al., 2009). Pseudowords were phonemically transcribed using the most common pronunciation.

## Acknowledgements

We express our gratitude to all the patients who participated in this study; the neurologists at the Texas Comprehensive Epilepsy Program who participated in the care of these patients; and the nurses and technicians in the Epilepsy Monitoring Unit at Memorial Hermann Hospital who helped make this research possible. This work was supported by the National Institute of Neurological Disorders and Stroke NS098981.

## Author Contributions

Conceptualization: OW, CD, SD, NT; Methodology: OW, CD, NT; Data curation: OW, CD, PSR, ZR; Software: OW, CD, AC; Formal Analysis: OW, AC; Writing – Original Draft: OW; Writing – Review and Editing: OW, SD, SFB, NT; Visualization: OW; Funding Acquisition: NT; Neurosurgical Procedures: NT.

## Supplementary Information

Video 1: **Spread of Stimulus-Locked Activity across the Cortical Surface**. MEMA movie of the time course of broadband gamma activation across the cortical surface with trials time-locked to onset of the visual stimulus. Regions in black did not have consistent coverage for reliable MEMA results.

Video 2: **Spread of Articulation-Locked Activity across the Cortical Surface**. MEMA movie of the time course of broadband gamma activation across the cortical surface with trials time locked to the onset of articulation. Regions in black did not have consistent coverage for reliable MEMA results.

**Supplementary Table 1:** Distribution of Statistical Regressors. Minimum, median and maximum values for each of the regressors used, across the whole stimulus set and for individual word classes. Statistical models used normalized data, subtracting the minimum value and dividing by the range across the whole stimulus set.

|  | All |  |  | Regular |  |  | Exception |  |  | Pseudowords |  |  |
| --- | --- | --- | --- | --- | --- | --- | --- | --- | --- | --- | --- | --- |
|  | Min | Med | Max | Min | Med | Max | Min | Med | Max | Min | Med | Max |
| **Length** | 3 | 4 | 6 | 3 | 4 | 6 | 3 | 4 | 6 | 4 | 4 | 6 |
| **Word Frequency** | -1 | 1.7 | 3.8 | -1 | 1.7 | 3 | -0.5 | 1.8 | 3.8 | - | - | - |
| **Orthographic Neighborhood** | 1 | 1.7 | 2.8 | 1 | 1.8 | 2.4 | 1 | 1.6 | 2 | 1.2 | 1.9 | 2.8 |
| **Phonological Neighborhood** | 0 | 22 | 49 | 6 | 24 | 42 | 1 | 22 | 49 | 0 | 19 | 41 |
| **Positional Letter Frequency** | 4.4 | 4.9 | 5.2 | 4.4 | 4.9 | 5.1 | 4.5 | 4.9 | 5.1 | 4.5 | 4.8 | 5.2 |

**Supplementary Figure 1:**
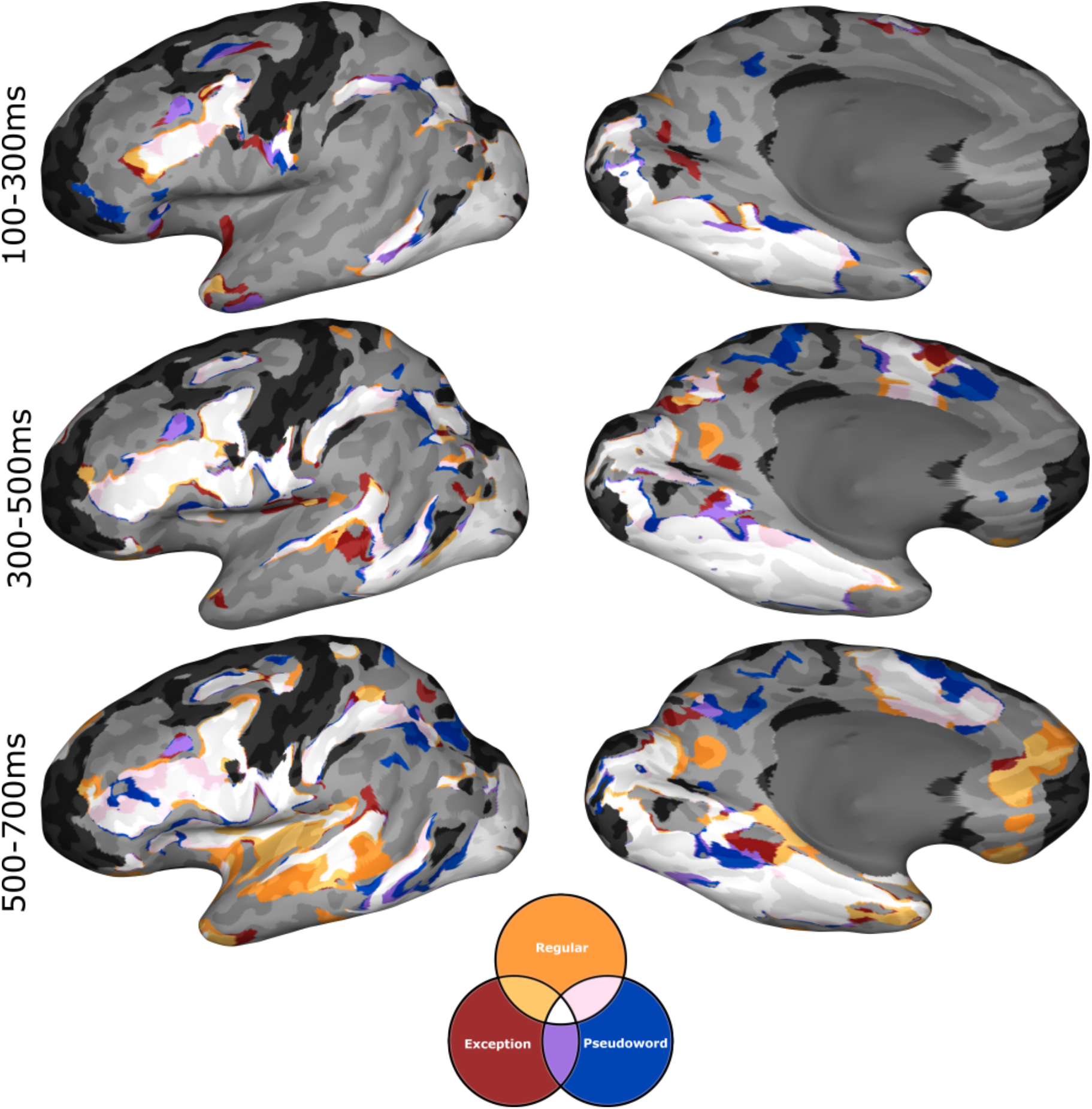
Conjunction Map of Word Class Activations. MEMA conjunction maps showing overlap of binarized activation maps of each of the three word classes tested (%BGA > 5%, *t* > 2.58, patients ≥ 3), over three time windows locked to stimulus onset. Across all time windows all three word classes demonstrate a gross overlap of activation (white). In the later time window, areas associated with post-articulatory processes (e.g. auditory cortex) show selective activation for known words, reflecting differences in response time between known words and novel pseudowords. Regions in black did not have consistent coverage for reliable MEMA results.

**Supplementary Figure 2:**
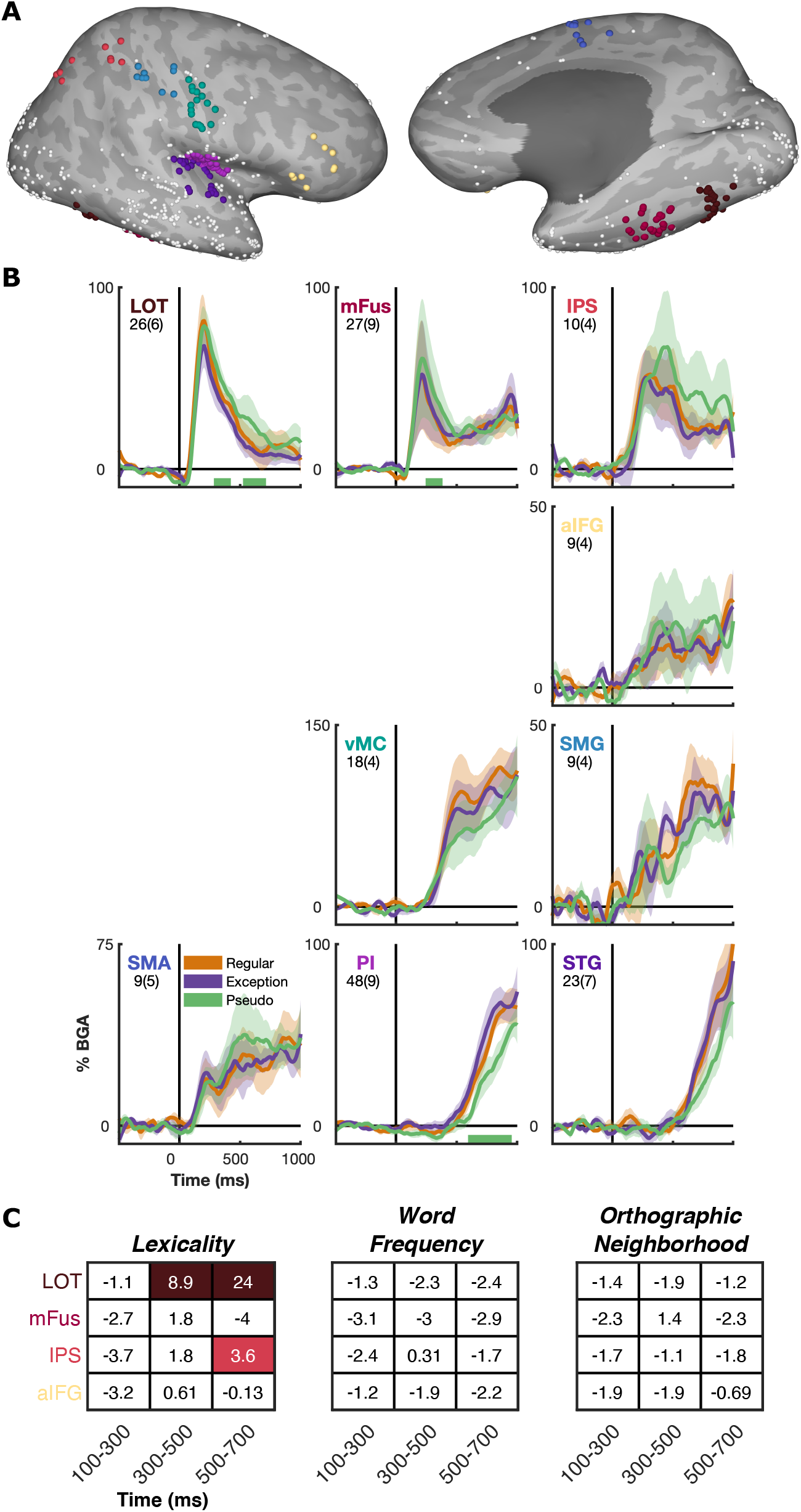
Spatiotemporal Activation Profiles in Right Hemisphere. (A) Right hemisphere analogs of the main analysis ROIs, highlighting all 588 responsive electrodes (out of 1,712 implanted in right hemisphere). (B) Mean activation (± SE) for each word class, within each ROI, during word reading, averaged within patient, time locked to stimulus onset. Number of electrodes and patients, per ROI, is indicated. Colored bars represent regions of significant difference from exception words (Wilcoxon signed-rank, p < 0.05 for >100ms). (C) Bayes factor analysis of lexicality, word frequency and orthographic neighborhood effects in the four pre-articulatory ROIs, for three time windows. Lexicality tested all known words against pseudowords. Word frequency was regressed across all known words. Orthographic neighborhood was regressed across all pseudowords. Bayes factor (ln(BF_10_)) shown for each contrast and values >2.3 are highlighted. LOT, Lateral OccipitoTemporal cortex; mFus, mid-Fusiform Cortex; IPS, Inferior Parietal Sulcus; aIFG, anterior Inferior Frontal Gyrus; vMC, ventral Motor Cortex; SMG, Supra Marginal Gyrus; SMA, Supplementary Motor Area; PI, Posterior Insula; STG, Superior Temporal Gyrus.

## References

Argall, B.D., Saad, Z.S., and Beauchamp, M.S. (2006). Simplified intersubject averaging on the cortical surface using SUMA. Hum. Brain Mapp. 27, 14–27.

Baldo, J. V, Wilkins, D.P., Ogar, J., Willock, S., and Dronkers, N.F. (2011). Role of the precentral gyrus of the insula in complex articulation. Cortex 47, 800–807.

Binder, J.R., Desai, R.H., Graves, W.W., and Conant, L.L. (2009). Where is the semantic system? A critical review and meta-analysis of 120 functional neuroimaging studies. Cereb. Cortex 19, 2767–2796.

Bouhali, F., Bézagu, Z., Dehaene, S., and Cohen, L. (2019). A mesial-to-lateral dissociation for orthographic processing in the visual cortex. Proc. Natl. Acad. Sci. U. S. A. 116, 21936–21946.

Brysbaert, M., and New, B. (2009). Moving beyond Kučera and Francis: A critical evaluation of current word frequency norms and the introduction of a new and improved word frequency measure for American English. Behav. Res. Methods 41, 977–990.

Cazzoli, D., Müri, R.M., Kennard, C., and Rosenthal, C.R. (2015). The Role of the Right Posterior Parietal Cortex in Letter Migration between Words. J. Cogn. Neurosci. 27, 377–386.

Cohen, L., Dehaene, S., Vinckier, F., Jobert, A., and Montavont, A. (2008). Reading normal and degraded words: Contribution of the dorsal and ventral visual pathways. Neuroimage 40, 353–366.

Coltheart, M., Rastle, K., Perry, C., Langdon, R., and Ziegler, J. (2001). DRC: A dual route cascaded model of visual word recognition and reading aloud. Psychol. Rev. 108, 204–256.

Conner, C.R., Chen, G., Pieters, T.A., and Tandon, N. (2014). Category specific spatial dissociations of parallel processes underlying visual naming. Cereb. Cortex 24, 2741–2750.

Cox, R.W. (1996). AFNI: Software for Analysis and Visualization of Functional Magnetic Resonance Neuroimages. Comput. Biomed. Res. 29, 162–173.

Dale, A.M., Fischl, B., and Sereno, M.I. (1999). Cortical Surface-Based Analysis: I. Segmentation and Surface Reconstruction. Neuroimage 9, 179–194.

Dehaene-Lambertz, G., Monzalvo, K., and Dehaene, S. (2018). The emergence of the visual word form: Longitudinal evolution of category-specific ventral visual areas during reading acquisition. PLOS Biol. 16, e2004103.

Dehaene, S., and Cohen, L. (2011). The unique role of the visual word form area in reading. Trends Cogn. Sci. 15, 254–262.

Dickens, J.V., Fama, M.E., DeMarco, A.T., Lacey, E.H., Friedman, R.B., and Turkeltaub, P.E. (2019). Localization of phonological and semantic contributions to reading. J. Neurosci. 39, 5361–5368.

Dronkers, N.F., Wilkins, D.P., Van Valin, R.D., Redfern, B.B., and Jaeger, J.J. (2004). Lesion analysis of the brain areas involved in language comprehension. Cognition 92, 145–177.

Esposito, F., Singer, N., Podlipsky, I., Fried, I., Hendler, T., and Goebel, R. (2013). Cortex-based inter-subject analysis of iEEG and fMRI data sets: Application to sustained task-related BOLD and gamma responses. Neuroimage 66, 457–468.

Fiebach, C.J., Friederici, A.D., Müller, K., and Von Cramon, D.Y. (2002). fMRI evidence for dual routes to the mental lexicon in visual word recognition. J. Cogn. Neurosci. 14, 11–23.

Fischl, B., Sereno, M.I., Tootell, R.B.H., and Dale, A. (1999). High-resolution inter-subject averaging and a surface-based coordinate system. Hum. Brain Mapp. 8, 272–284.

Fridriksson, J., Den Ouden, D.B., Hillis, A.E., Hickok, G., Rorden, C., Basilakos, A., Yourganov, G., and Bonilha, L. (2018). Anatomy of aphasia revisited. Brain 141, 848–862.

Glasser, M.F., Coalson, T.S., Robinson, E.C., Hacker, C.D., Harwell, J., Yacoub, E., Ugurbil, K., Andersson, J., Beckmann, C.F., Jenkinson, M., et al. (2016). A multi-modal parcellation of human cerebral cortex. Nature 536, 171–178.

Glezer, L.S., Kim, J., Rule, J., Jiang, X., and Riesenhuber, M. (2015). Adding Words to the Brain’s Visual Dictionary: Novel Word Learning Selectively Sharpens Orthographic Representations in the VWFA. J. Neurosci. 35, 4965–4972.

Gramfort, A. (2013). MEG and EEG data analysis with MNE-Python. Front. Neurosci. 7, 1–13.

Gramfort, A., Luessi, M., Larson, E., Engemann, D.A., Strohmeier, D., Brodbeck, C., Parkkonen, L., and Hämäläinen, M.S. (2014). MNE software for processing MEG and EEG data. Neuroimage 86, 446–460.

Graves, W.W., Desai, R., Humphries, C., Seidenberg, M.S., and Binder, J.R. (2010). Neural systems for reading aloud: A multiparametric approach. Cereb. Cortex 20, 1799–1815.

Hartwigsen, G., Baumgaertner, A., Price, C.J., Koehnke, M., Ulmer, S., and Siebner, H.R. (2010). Phonological decisions require both the left and right supramarginal gyri. Proc. Natl. Acad. Sci. U. S. A. 107, 16494–16499.

Heim, S., Wehnelt, A., Grande, M., Huber, W., and Amunts, K. (2013). Effects of lexicality and word frequency on brain activation in dyslexic readers. Brain Lang. 125, 194–202.

Hirshorn, E.A., Li, Y., Ward, M.J., Richardson, R.M., Fiez, J.A., and Ghuman, A.S. (2016). Decoding and disrupting left midfusiform gyrus activity during word reading. Proc. Natl. Acad. Sci. 113, 8162–8167.

Hula, W.D., Panesar, S., Gravier, M.L., Yeh, F.-C., Dresang, H.C., Dickey, M.W., and Fernandez-Miranda, J.C. (2020). Structural white matter connectometry of word production in aphasia: an observational study. Brain 143, 2532–2544.

Jobard, G., Crivello, F., and Tzourio-Mazoyer, N. (2003). Evaluation of the dual route theory of reading: A metanalysis of 35 neuroimaging studies. Neuroimage 20, 693–712.

Kadipasaoglu, C.M., Baboyan, V.G., Conner, C.R., Chen, G., Saad, Z.S., and Tandon, N. (2014). Surface-based mixed effects multilevel analysis of grouped human electrocorticography. Neuroimage 101, 215–224.

Kaestner, E., Wu, X., Friedman, D., Dugan, P., Devinsky, O., Carlson, C., Doyle, W., Thesen, T., and Halgren, E. (2021). The Precentral Gyrus Contributions to the Early Time-Course of Grapheme-to-Phoneme Conversion. Neurobiol. Lang. 1–61.

Kleiner, M., Brainard, D., and Pelli, D. (2007). What’s new in Psychtoolbox-3? Perception 36.

Kronbichler, M., Hutzler, F., Wimmer, H., Mair, A., Staffen, W., and Ladurner, G. (2004). The visual word form area and the frequency with which words are encountered: Evidence from a parametric fMRI study. Neuroimage 21, 946–953.

Liu, Y., Shi, G., Li, M., Xing, H., Song, Y., Xiao, L., Guan, Y., and Han, Z. (2021). Early top-down modulation in visual word form processing: Evidence from an intracranial SEEG study. J. Neurosci. 41, JN-RM-2288-20.

Lochy, A., Jacques, C., Maillard, L., Colnat-Coulbois, S., Rossion, B., and Jonas, J. (2018). Selective visual representation of letters and words in the left ventral occipito-temporal cortex with intracerebral recordings. Proc. Natl. Acad. Sci. 115, E7595–E7604.

Mălîia, M.-D., Donos, C., Barborica, A., Popa, I., Ciurea, J., Cinatti, S., and Mîndru, I. (2018). Functional mapping and effective connectivity of the human operculum. Cortex 109, 303–321.

Martin, A., Schurz, M., Kronbichler, M., and Richlan, F. (2015). Reading in the brain of children and adults: A meta-analysis of 40 functional magnetic resonance imaging studies. Hum. Brain Mapp. 36, 1963–1981.

Matchin, W., and Hickok, G. (2020). The Cortical Organization of Syntax. Cereb. Cortex 30, 1481–1498.

Matchin, W., Hammerly, C., and Lau, E. (2017). The role of the IFG and pSTS in syntactic prediction: Evidence from a parametric study of hierarchical structure in fMRI. Cortex 88, 106–123.

Nobre, A.C., Allison, T., and McCarthy, G. (1994). Word recognition in the human inferior temporal lobe. Nature 372, 260–263.

Numssen, O., Bzdok, D., and Hartwigsen, G. (2021). Functional specialization within the inferior parietal lobes across cognitive domains. Elife 10.

Perry, C., Ziegler, J.C., and Zorzi, M. (2007). Nested incremental modeling in the development of computational theories: The CDP+ model of reading aloud. Psychol. Rev. 114, 273–315.

Perry, C., Ziegler, J.C., and Zorzi, M. (2010). Beyond single syllables: Large-scale modeling of reading aloud with the Connectionist Dual Process (CDP++) model. Cogn. Psychol. 61, 106–151.

Perry, C., Zorzi, M., and Ziegler, J.C. (2019). Understanding Dyslexia Through Personalized Large-Scale Computational Models. Psychol. Sci. 30, 386–395.

Pieters, T.A., Conner, C.R., and Tandon, N. (2013). Recursive grid partitioning on a cortical surface model: an optimized technique for the localization of implanted subdural electrodes. J. Neurosurg. 118, 1086–1097.

Price, C.J., and Devlin, J.T. (2011). The Interactive Account of ventral occipitotemporal contributions to reading. Trends Cogn. Sci. 15, 246–253.

Rapcsak, S.Z., Beeson, P.M., Henry, M.L., Leyden, A., Kim, E., Rising, K., Andersen, S., and Cho, H.S. (2009). Phonological dyslexia and dysgraphia: Cognitive mechanisms and neural substrates. Cortex 45, 575–591.

Rapp, B., Purcell, J., Hillis, A.E., Capasso, R., and Miceli, G. (2016). Neural bases of orthographic long-term memory and working memory in dysgraphia. Brain 139, 588–604.

Raschle, N.M., Chang, M., and Gaab, N. (2011). Structural brain alterations associated with dyslexia predate reading onset. Neuroimage 57, 742–749.

Ripamonti, E., Aggujaro, S., Molteni, F., Zonca, G., Frustaci, M., and Luzzatti, C. (2014). The anatomical foundations of acquired reading disorders: A neuropsychological verification of the dual-route model of reading. Brain Lang. 134, 44–67.

Rollo, P.S., Rollo, M.J., Zhu, P., Woolnough, O., and Tandon, N. (2020). Oblique trajectory angles in robotic stereo-electroencephalography. J. Neurosurg.

Saad, Z.S., and Reynolds, R.C. (2012). Suma. Neuroimage 62, 768–773.

Sebastian, R., Gomez, Y., Leigh, R., Davis, C., Newhart, M., and Hillis, A.E. (2014). The roles of occipitotemporal cortex in reading, spelling, and naming. Cogn. Neuropsychol. 31, 511–528.

Shim, H.S., Hurley, R.S., Rogalski, E., and Mesulam, M.M. (2012). Anatomic, clinical, and neuropsychological correlates of spelling errors in primary progressive aphasia. Neuropsychologia 50, 1929–1935.

Simos, P.G., Breier, J.I., Fletcher, J.M., Foorman, B.R., Castillo, E.M., and Papanicolaou, A.C. (2002). Brain mechanisms for reading words and pseudowords: An integrated approach. Cereb. Cortex 12, 297–305.

Sliwinska, M.W., James, A., and Devlin, J.T. (2015). Inferior Parietal Lobule Contributions to Visual Word Recognition. J. Cogn. Neurosci. 27, 593–604.

Stoeckel, C., Gough, P.M., Watkins, K.E., and Devlin, J.T. (2009). Supramarginal gyrus involvement in visual word recognition. Cortex 45, 1091–1096.

Tandon, N. (2012). Mapping of human language. In Clinical Brain Mapping, D. Yoshor, and E. Mizrahi, eds. (McGraw Hill Education), pp. 203–218.

Tandon, N., Tong, B.A., Friedman, E.R., Johnson, J.A., Von Allmen, G., Thomas, M.S., Hope, O.A., Kalamangalam, G.P., Slater, J.D., and Thompson, S.A. (2019). Analysis of Morbidity and Outcomes Associated With Use of Subdural Grids vs Stereoelectroencephalography in Patients With Intractable Epilepsy. JAMA Neurol. 76, 672–681.

Taylor, J.S.H., Rastle, K., and Davis, M.H. (2013). Can cognitive models explain brain activation during word and pseudoword reading? A meta-analysis of 36 neuroimaging studies. Psychol. Bull. 139, 766–791.

Taylor, J.S.H., Rastle, K., and Davis, M.H. (2014). Interpreting response time effects in functional imaging studies. Neuroimage 99, 419–433.

Taylor, J.S.H., Davis, M.H., and Rastle, K. (2019). Mapping visual symbols onto spoken language along the ventral visual stream. Proc. Natl. Acad. Sci. 116, 17723–17728.

Temple, E., Deutsch, G.K., Poldrack, R.A., Miller, S.L., Tallal, P., Merzenich, M.M., and Gabrieli, J.D.E. (2003). Neural deficits in children with dyslexia ameliorated by behavioral remediation: Evidence from functional MRI. Proc. Natl. Acad. Sci. U. S. A. 100, 2860–2865.

Tomasino, B., Ius, T., Skrap, M., and Luzzatti, C. (2020). Phonological and surface dyslexia in individuals with brain tumors: Performance pre-, intra-, immediately post-surgery and at follow-up. Hum. Brain Mapp. 41, 5015–5031.

Tong, B.A., Esquenazi, Y., Johnson, J., Zhu, P., and Tandon, N. (2020). The Brain is Not Flat: Conformal Electrode Arrays Diminish Complications of Subdural Electrode Implantation, A Series of 117 Cases. World Neurosurg. 144, e734–e742.

Tourville, J.A., and Guenther, F.H. (2011). The DIVA model: A neural theory of speech acquisition and production. Lang. Cogn. Process. 26, 952–981.

Vaden, K.I., Halpin, H.R., and Hickok, G.S. (2009). Irvine Phonotactic Online Dictionary, Version 2.0. [Data file].

Vinckier, F., Naccache, L., Papeix, C., Forget, J., Hahn-Barma, V., Dehaene, S., and Cohen, L. (2006). “What” and “where” in word reading: Ventral coding of written words revealed by parietal atrophy. J. Cogn. Neurosci. 18, 1998–2012.

Whaley, M.L., Kadipasaoglu, C.M., Cox, S.J., and Tandon, N. (2016). Modulation of Orthographic Decoding by Frontal Cortex. J. Neurosci. 36, 1173–1184.

White, A.L., Palmer, J., Boynton, G.M., and Yeatman, J.D. (2019). Parallel spatial channels converge at a bottleneck in anterior word-selective cortex. Proc. Natl. Acad. Sci. 116, 10087–10096.

Wilson, S.M., DeMarco, A.T., Henry, M.L., Gesierich, B., Babiak, M., Mandelli, M.L., Miller, B.L., and Gorno-Tempini, M.L. (2014). What Role Does the Anterior Temporal Lobe Play in Sentence-level Processing? Neural Correlates of Syntactic Processing in Semantic Variant Primary Progressive Aphasia. J. Cogn. Neurosci. 26, 970–985.

Woodhead, Z.V.J., Barnes, G.R., Penny, W., Moran, R., Teki, S., Price, C.J., and Leff, A.P. (2014). Reading front to back: MEG evidence for early feedback effects during word recognition. Cereb. Cortex 24, 817–825.

Woolnough, O., Forseth, K.J., Rollo, P.S., and Tandon, N. (2019). Uncovering the functional anatomy of the human insula during speech. Elife 8, e53086.

Woolnough, O., Donos, C., Rollo, P.S., Forseth, K.J., Lakretz, Y., Crone, N.E., Fischer-Baum, S., Dehaene, S., and Tandon, N. (2021). Spatiotemporal dynamics of orthographic and lexical processing in the ventral visual pathway. Nat. Hum. Behav. 5, 389–398.

Xu, W., Kolozsvari, O.B., Oostenveld, R., and Hämäläinen, J.A. (2020). Rapid changes in brain activity during learning of grapheme-phoneme associations in adults. Neuroimage 220, 117058.

Yarkoni, T., Balota, D., and Yap, M. (2008). Moving beyond Coltheart’s N: A new measure of orthographic similarity. Psychon. Bull. Rev. 15, 971–979.

Zhi, D., King, M., and Diedrichsen, J. (2021). Evaluating brain parcellations using the distance controlled boundary coefficient. BioRxiv 2021.05.11.443151.

